# *In vitro* and *in silico* analyses of the angiotensin-I converting enzyme inhibitory activity of peptides identified from *Bellamya bengalensis* protein hydrolysates

**DOI:** 10.1101/2020.04.09.034306

**Authors:** Tanmoy Kumar Dey, Roshni Chatterjee, Anadi Roychoudhury, Debjyoti Paul, Rahul Shubhra Mandal, Souvik Roy, Pubali Dhar

**Affiliations:** Laboratory of Food Science and Technology, Food and Nutrition Division, University of Calcutta, 20B, Judges Court Road, Alipore, Kolkata, West Bengal, India, PIN: 700027; Centre for Nanoscience and Nanotechnology, University of Calcutta; Department of Physiology, Serampur College (Autonomous), University of Calcutta, 8, William Carey Sarani, Maniktala, Serampore, West Bengal, India, PIN: 712201; Biomedical Informatic Centre, National Institute of Cholera and Enteric Diseases, Scheme XM, P-33, CIT Road, Beliaghata, Kolkata, West Bengal, India. PIN: 700010; DBT-Interdisciplinary Programme of Life Sciences (DBT-IPLS), Modern Biology wing, University of Calcutta, 35, Ballygunge Circular Road, Kolkata, West Bengal, India. PIN: 700019

**Author notes:** Corresponding Author: Dr. Pubali Dhar Associate Professor, Laboratory of Food Science and Technology, Food and Nutrition Division, University of Calcutta, 20B, Judges Court Road, Alipore, Kolkata, West Bengal, India, PIN: 700027.

**Keywords:** *Bellamya bengalensis* protein hydrolysate, Alcalse, ACE-inhibitory activity, Lisinopril, Isothermal Titration Calorimetry, uncompetitive inhibition, co-operative ligand binding, site-specific molecular docking

## Abstract

The study focuses on identification of ACE-inhibitory peptides from the proteolytic digests of muscle protein of *Bellamya bengalensis* and its underlying mechanism. 120 min Alcalase-hydrolysates were ultrafiltered to isolate the small peptide fraction (<3kDa) and *in vitro* ACE-inhibitory activity was analyzed. The IC_50_ value of the 120 min hydrolysate ultafiltered fraction was found to be 86.74 ± 0.575 µg/mL, while the IC_50_ of Lisinopril is 0.31 ± 0.07 µg/mL. This fraction was assessed in MALDI-ToF mass-spectrometer and five peptides were sequenced via *de novo* sequencing. The ACE-inhibitory potential of the peptides have a positive correlation with the hydrophobicity of the amino acids. Synthetic analogue of the peptide (IC_50_ value 8.52 ± 0.779 µg/mL) was used to understand the thermodynamics of the inhibition by checking the binding affinity of the peptide to ACE by Isothermal titration calorimetry compared with lisinopril, and further substantiated by *in silico* site specific molecular docking study.

## 1. Introduction

A number of naturally grown traditional food items are known to have precise therapeutic roles against specific disease aetiology (Kwon, Apostolidis, Kim, & Shetty, 2007). Current foodomic research has identified the effective ‘nutraceuticals’ and established their health-promoting activities of such traditional food items, but most of these traditional food items, present in the nature, still remain to be analyzed for their health promoting attributes. Incorporation of natural nutraceuticals are much better alternatives than synthetic pharmaceuticals as the xenobiotics can’t exactly mimic natural behavior of the biomolecules within the physiological microenvironment (Bhattacharya et al., 2014). Hence the modern concept of diet and nutrition focuses on making our daily foodstuffs more enriched or more fortified with such natural nutraceuticals to make them more healthy and ‘functional’ in terms of boosting and maintaining health parameters.

Fresh water molluscs are one such traditional food item which serves as an extremely appreciated cuisine in European countries like Netherlands, France, Austria etc. since long time. In the Indian context, these fresh water molluscs, especially the gastropod snail *Bellamya bengalensis* provides a major source of animal protein, mostly among the tribal population of both India and Bangladesh (Prabhakar & Roy 2009; Baby, Hasan, Kabir, & Naser, 2010).

World-over, there is a promising trend towards better utilization of small ‘bioactive peptides’ for their important nutraceutical potential, i.e. antioxidative, antihypertensive, anti-cancer etc. (Udenigwe, & Aluko, 2012). Bioactive peptides with 2-10 amino acids, isolated from natural food components by enzymatic bioprocessing techniques, are conceptualized to function better within body due to better digestibility and higher bioavailability, although scientific ratification of the molecular mechanism is still inadequate. Among various bioactive peptides that have been studied extensively, antihypertensive peptides need special mention. Enzymatic hydrolysis of numerous sources like fish proteins, seed proteins especially from legumes and oilseeds, milk proteins etc. have been studied for short peptides with potent Angiotensin I converting enzyme (ACE) inhibitory activity (Udenigwe, & Aluko, 2012; Aluko, 2015). The ACE inhibitors play a pivotal role in the regulation of blood pressure. It acts through the renin–angiotensin–aldosterone system (RAAS) to control blood pressure of the body. ACE (E.C. 3.4.15.1.), a dipeptidylcarboxypeptidase plays a key role in the RAAS system by activating vasoconstrictor angiotensin-II by cleaving His-Leu dipeptide from the angiotensin I (Asp-Arg-Val-Tyr-Ile-His-Pro-Phe-His-Leu) (Natesh, Schwger, Sturrock, & Acharya, 2003) and inactivating vasodilator bradykinin (Raghavan & Kristinsson, 2009), thereby reducing blood pressure and maintaining electrolyte equilibrium in blood. Successful inhibition of ACE can thus regulate hypertension. Synthetic ACE inhibitors like captopril, lisinopril, enalpril etc. are being used as effective measure for treating hypertension but their utilization causes countless side effects like allergy, dermatitis, cough, etc. (Chen, Wang, Ye, Wu, & Xia, 2013). Herein lays the importance of nutraceuticals as natural alternative ACE inhibitors with no such adverse effects.

The objective of the present work was to explore the bioactive short peptides from enzymatic digests of *Bellamya bengalensis* protein isolate for both nutritional and therapeutic benefits with targeted action. Further emphasis was laid on understanding the mechanism of the biochemical interactions between the bioactive short peptides and ACE. The findings of the work would reveal the translational potential of such low cost highly abundant, non-conventional food resource for biomedical application.

## 2. Materials and Methods

### 2.1. Collection of materials

*Bellamya bengalensis* f. *typica (Lamark 1822)* samples were collected from freshwater water bodies of Kakdweep, South 24 Parganas, West Bengal, eastern part of India and as identified by Zoological Survey of India, Prani Vigyan Bhaban, M Block, New Alipore, Kolkata-53. [F. No. 229–10/98-Mal./7606]. After removing the shells, the foot muscle (edible snail meat) was separated from viscera, washed in clean water and minced before storing in −80°C.

### 2.2. Preparation of B. bengalensis protein concentrates (BBPC)

*B. bengalensis* meat was homogenized (Ultra Turrax T18, IKA®Werke GmbH & Co., KG, Stufen, Germany) in phosphate buffer (pH 7.0) at a ratio of 1:10 for 60 min at 12000 rpm. The homogenate was centrifuged and the protein-rich supernatant was collected by partitioning over chloroform. The protein-rich fraction was lyophilized under partial vacuum upto dry powder form and stored at - 80°C untill further use.

### 2.5. Preparation of B. bengalensis protein hydrolysate (BBPH)

15 g/L of BBPC was hydrolyzed by Alcalase®2.4 L (from *Bacillus licheniformis*, Subtilisin A, P4860, Sigma Aldrich, MO, USA) with different enzyme concentration (0.1% v/v to 0.5% v/v) at conditions as optimized by Ahn, Jeon, Kim, & Je, 2012. The reaction was stopped at time intervals i.e. 0 min, 10 min, 30 min, 60 min, 90 min and 120 min, by inactivation of alcalase (Mohammad, Kumar, & Basha, 2015) and the hydrolysates were stored at −20ºC until further use (Chatterjee, Dey, Ghosh, & Dhar, 2015).

Degree of hydrolysis (DH; %) was determined by Ninhydrin method (Molnar-Perl & Pinter-Szakacs, 1989). DH was defined as the percentage of peptide bonds hydrolyzed was calculated by the determination of free amino groups reacting with Ninhydrin. The degree of hydrolysis was determined from the following equation:

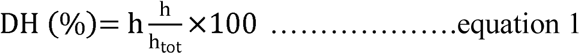

where, h = concentration of peptide bond hydrolysed (meq/gm) and h_tot_ = total amount of peptide bond, taken as 8 amino meq/gm.

*B. bengalensis* protein hydrolysates (BBPH) were subjected to centrifugal ultrafiltration using 3kDa cut-off membrane filtration unit (Vivaspin20, VS2091, Sartorius AG, Goettingen, Germany) to separate the low molecular peptide fraction (< 3kDa). The ultrafiltered samples were lyophilized under partial vacuum until dry and stored as fine powder in −80℃ until they were further used for bioactivity assays.

### 2.6. In vitro ACE inhibitory assay

*In vitro* ACE inhibitory activity assay was performed on the basis of the method as described by Li, Liu, Shi, & Le, 2005 with slight modifications. 200 µl of ACE (20 mU/mL; from Rabbit lung, A67778, Sigma-Aldrich, MO, USA) was added to 2.17mM hippuryl-L-histidyl-L-leucine (HHL, H1635, Sigma-Aldrich, MO, USA) in 100mM sodium borate buffer (pH 8.3) containing 300mM NaCl to start the reaction by incubating at 37°C for 30 min. 50 µL of 1 mg/mL (w/v) short peptides (< 3kDa, ultrafiltered) in sodium borate buffer, isolated from the BBPHs were added to the reaction mix before adding the enzyme to assess the inhibition potential of the peptides and the results have been compared with a standard ACE inhibitor Lisinopril (L2777, Sigma-Aldrich, MO, USA), used as positive control (50 µL of 2 µg/mL).

The reaction was stopped by 125 µL of 1M HCl and 30 sec later the reaction was neutralized by adding 0.5 mL of 1M NaOH. 2 mL of diluent buffer containing 0.2M KH_2_PO_4_ and 1.5 mL of the colour reagent containing cyanuric chloride in 1, 4-dioxane was added, vortexed and centrifuged to remove any particulate matter. In order to evaluate the *in vitro* activity of ACE, the chromogen developed by the reaction of cyanuric chloride with the released hippurate, was measured spectrophotometrically at 382 nm against corresponding reagent blank. The ACE inhibition activity was calculated using the following equation:

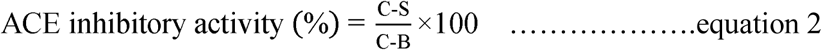

where, C is the optical density of the control reaction mixture consisting of ACE and substrate HHL; S is the optical density of the reaction mixture in presence of the peptide sample and B is the optical density of blank.

For every sample, a gradient of concentration of the sample were added to the reaction for effective inhibition of ACE. The ACE inhibitory potential of each sample were expressed in terms of their IC_50_ value, defined as the concentration of the sample required to inhibit 50% of the ACE activity.

### 2.7. Identification and preparation of ACE inhibitory peptide

Based on best ACE-inhibitory activity, the sub-3kDa fraction of the 120 minute hydrolysate (BBPH_A120_) was analyzed by matrix-assisted laser desorption/ionization time-of-flight (MALDI-ToF) mass spectrometry (Chatterjee et al., 2015) for identifying peptides probably responsible for the inhibitory activity, using Ultraflextreme ToF/ToF mass spectrometer (Bruker Daltonics, Bremen, Germany). The Mass Spectra (600-3000 *m/z*) were acquired in reflector mode and from the mass spectrum, the signals with particularly high intensity (> 1 a.u.) were further analyzed as their MS-MS spectra were acquired in LIFT mode using Flex Control (version 3.4) software, with 2000 shots added per sample. From the MS-MS spectra of the selected peptides, to *de novo* was sequencing to identify the amino acid sequence of the peptides.

These peptide sequences were cross-matched with the sequences enlisted in the AHTPDB database (http://crdd.osdd.net/raghava/ahtpdb/) of the established ACE inhibitory peptides isolated from edible sources reported till date. The peptide sequence (henceforth referred as Belpep) with highest match (mostly with previously reported peptides isolated from milk protein, soy protein and pea protein; especially with low IC_50_ values), was chosen for synthesized by solid-phase method, using standard Fmoc-chemistry, at Pepmic Solutions, Suzhou, China. The purity (95-99.9%) of the synthesized peptide was confirmed by high-performance liquid chromatography (HPLC) and the molecular weight was verified by mass spectrometry (MS) study. The pure peptide was utilized to validate its ACE inhibitory activity by assessing the hippuric acid release adding 50 µL of Belpep (2 µg/mL) in sodium borate buffer as mentioned above. This activity is further mechanistically validated for enzyme-substrate binding via isothermal calorimetric (ITC) analysis and the *in silico* modelling, using lisinopril as positive control.

### 2.8. Determination of the kinetics of ACE inhibitory activity of the peptide

The kinetics of the reaction catalyzed by ACE where HHL was hydrolysed to release the hippuric acid, was studied both in presence and absence of inhibitor, i.e. belpep as described in the present study. ACE activity was measured at various substrate (HHL) concentrations i.e. 0.5mM, 1.0mM, 2.0mM and 4.0mM, while the enzyme concentration was maintained at 20 mU/mL. The concentration of the inhibitor belpep used in the study was 1.0 mg/mL. The kinetic parameters (V_max_, K_m_) of the enzyme kinetics were established through the Lineweaver-Burk plot, in presence or absence of inhibitor. The type of the mechanism of belpep mediated inhibition, whether competitive, noncompetitive or uncompetitive, was also interpreted from the Lineweaver-Burk plot (Ahn et al., 2012).

### 2.9. Isothermal Titration Calorimetry

The Isothermal Titration Calorimetry (ITC) experiments were performed using Isothermal Titration Calorimeter (MicroCal iTC200, GE Healthcare Bioscience limited). Enzyme-substrate reactions were carried out within the reaction cell at 37°C, with a sodium borate buffer (pH 8.3) background. In a control experiment, the reaction cell was loaded with 350 µL ACE (0.357 µmol/L) which was titrated with 20 identical injections of 2 µL of HHL (1.0 mmol/L). The time interval between two injections was 120 sec. The heat change in the reaction cell due to the enzyme-substrate reaction was measured against a thermally stabilized reference cell. The thermogram peaks corresponding to the heat change in the reaction cell were integrated using ORIGIN 6 software (Microcal) supplied with the instrument (Ni, Li, Liu, & Hu, 2012). In the negative control experiment, only the buffer was injected in the reaction cell, without the enzyme. In the reactions involving inhibitors, the ITC cell was filled with 350 µL of ACE (0.357 µmol/L sodium borate buffer, pH 8.3) and 50 µL of pure belpep (0.15 mg/mL in sodium borate buffer, pH 8.3), which was then titrated with HHL (1.0 mmol/L) as described earlier. The thermograms were compared to assess the binding mechanism of the belpep.

### 2.10. Molecular Docking studies

The model for ACE used in this study was imported from the Protein Data Bank (1O86.pdb) which represented the crystal structure of the human angiotensin converting enzyme-lisinopril complex at 2 Å resolution (Natesh et al., 2003). The co-crystal structure as the basic mold was chosen because lisinopril is known to be one of the competitive inhibitor of the enzyme; thus localization of lisinopril within the ACE structure would identify its main catalytic site. ChemOffice 2004 software (CambridgeSoft Co., MA, USA) was used to construct the initial structure of the purified peptide. Before the docking between the belpep sequence and ACE, the cofactors like zinc and chloride atoms were retained in the active site, fixed to their crystal positions in the ACE-lisinopril complex (1O86.pdb) whereas, the lisinopril and the water molecules were removed.The automated molecular docking of the peptide with Human Angiotensin Converting Enzyme (ACE) was performed with through GalaxyPepDock web server (Lee, Heo, Lee, & Seok, 2015) using all default parameters. The binding site was selected as described by Jimsheena & Gowda, 2010, so that the binding site of lisinopril within the crystal structure of ACE was well sampled with a grid resolution of 0.3 Å. The two- and three-dimensional structures of the resulted docked structures were visualized and analyzed in PyMol software. The best docking pose of the belpep in the active site of ACE was obtained on the basis of least binding energy value and further analyzed to identify the hydrogen bonds and other hydrophobic or hydrophilic interactions between the amino acid residues at the ACE active site and the belpep.

### 2.11. Data analysis

All results were presented as means ± standard deviation (SD) of a minimum of 3–5 replicates of data. Data comparisons and analyses were done using the software Origin 8.1 (OriginLab Corporation, MA, USA). Significance of differences between two samples was determinedby the Student t-test. Significance limit was set at *p* < 0.05.

## 3. Results and Discussion

### 3.1. Degree of hydrolysis of B. bengalensis protein concentrate by Alcalase and their ACE inhibitory activities

The alcalase mediated hydrolysis curve of *Bellamya bengalensis* protein concentrate (BBPC) along a time gradient of 120 min was shown in **Supplementary Figure 1**. It is evident that with the increase in enzyme concentration (% v/v), degree of hydrolysis were increasing until it reached an optimum concentration (0.3% v/v). A highest degree of hydrolysis of 67.43 % was achieved at 120 min with 0.3% (v/v) enzyme concentration at a substrate concentration as high as 15 g/L. Demirhan, Apar, & Özbek, 2011 also reported an optimum hydrolytic activity at similar conditions. Further increase in the enzyme concentration didn’t showed significant better hydrolytic activity. Also extending the reaction beyond 120 min, the increase in degree of hydrolysis was found to be insignificant. Similar results were reported by Bhaskar, Benila, Radha, & Lalitha, 2008, where the *Catla catla* visceral protein was utilized as substrate. The increase in degree of hydrolysis was inferred as the increase in the number of cleaved peptide bonds and amplification in the number of smaller peptides.

**FIGURE 1:**
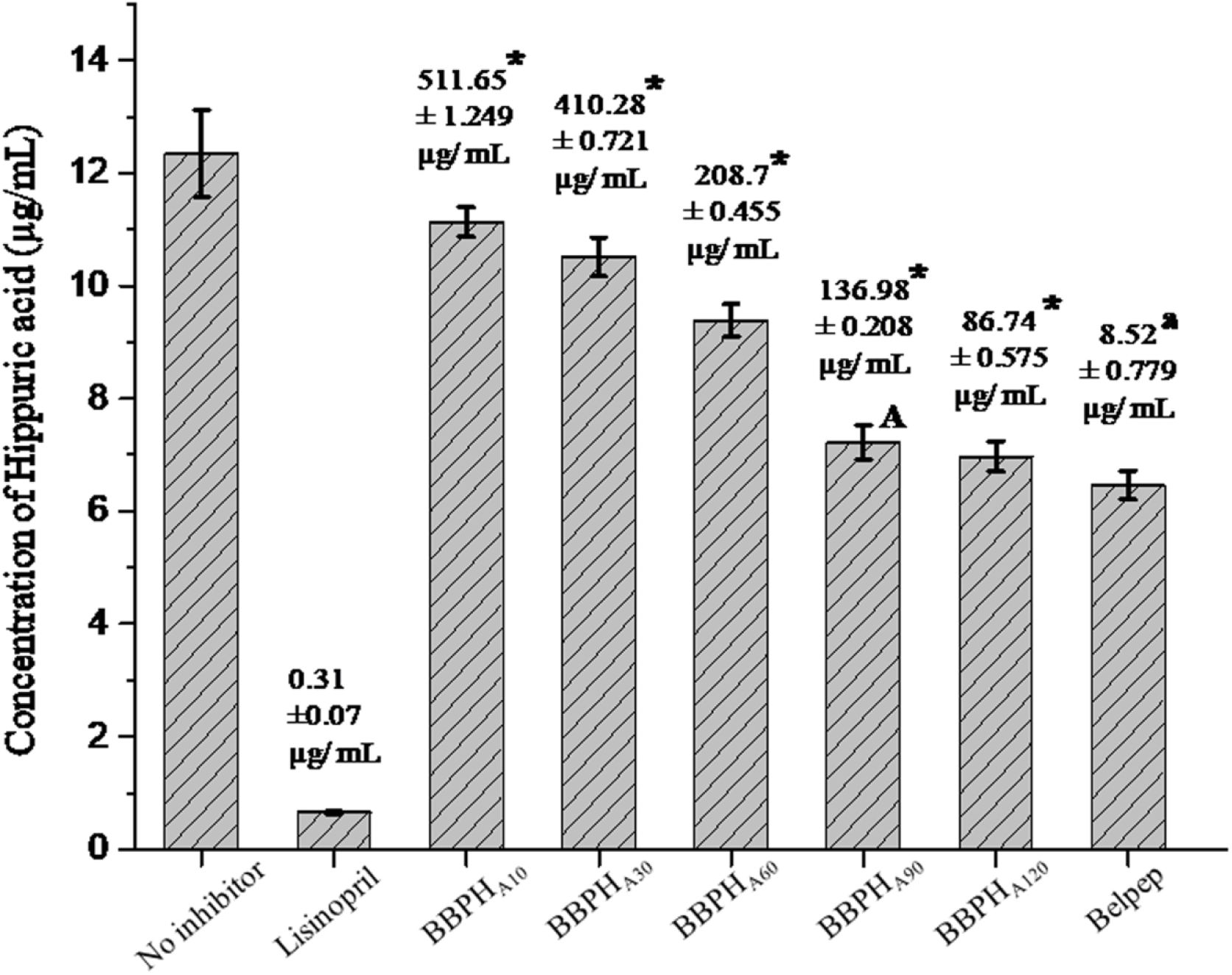
ACE INHIBITORY ACTIVITY (AS SHOWN BY HIPPURIC ACID RELEASE FROM HHL BY ACE) OF STANDARD ACE-INHIBITOR LISINOPRIL (AT A DOSE OF 50 µL OF 2 µG/ML SOLUTION) AND *B. BENGALENSIS* PROTEIN HYDROLYSATES (BBPHS, AT A DOSE OF 50 µL OF 1MG/ML SOLUTION) OF DIFFERENT TIME GRADIENT E.G. 10 MIN (BBPH_A10_), 30 MIN (BBPH_A30_), 60 MIN (BBPH_A60_), 90 MIN (BBPH_A90_) AND 120 MIN (BBPH_A120_) RESPECTIVELY AND THE SYNTHESIZED PEPTIDE, MENTIONED AS BELPEP (AT A DOSE OF 50 µL OF 0.15 MG/ML SOLUTION). IC_50_ VALUES OF EACH REACTION SET HAS BEEN SHOWN OVER EACH BAR AS MEAN ± SD VALUE, N=3. THE CAPITAL LETTER SUPERSCRIPT LETTER INDICATE THE SIGNIFICANT REDUCTION OF HIPPURATE RELEASE (*P* < 0.05), AS AFFECTED BY BBPH_A90_, IN COMPARISON TO BBPH_A60_ (*P* = 0.01987). THE * SIGNIFY THE SIGNIFICANT DIFFERENCE BETWEEN THE IC_50_ VALUES OF HYDROLYSATES OF TWO SUBSEQUENT TIME POINTS [BBPH_A10_ VS BBPH_A30_ (*P* = 0.0000139); BBPH_A30_ VS BBPH_A60_ (*P* = 0.0000029); BBPH_A60_ VS BBPH_A90_ (*P* = 0.0000075); BBPH_A90_ VS BBPH_A120_ (*P* = 0.0000211)]. THE SMALL LETTER SUPERSCRIPT SIGNIFY THE SIGNIFICANT DIFFERENCE IN THE IC_50_ VALUES OF BELPEP AND BBPH_A120_ (*P* = 0.0000992).

The pertinent literature suggests that smaller peptides with low molecular weight have shown better ACE-inhibitory potential (Chalé, Ruiz, Fernández, Ancona, & Campos, 2014; Wu, Jia, Yan, Du, & Gui, 2015; Wua, Du, Jia, & Kang, 2016). Hence, peptide fractions <3kDa were fractionated using an ultrafiltration membrane (Vivaspin®20, Sartorius, India). The filtrates were lyophilized and further solubilized in sterile buffered medium to assess their ACE-inhibitory activity and compared with Lisinopril (Figure 1). Being the standard inhibitor, Lisinopril reduced hippuric acid release (0.65 ± 0.03 µg/mL) significantly (*p* = 0.00154) compared to the uninhibited ACE activity (12.34 ± 0.774 µg/mL). Among all the alcalase hydrolysate groups, hydolysates at 120 min (BBPH_A120_, *Bellamya bengalensis* protein hydrolysates by Alcalse for 120 min) had shown the maximum inhibitory effect, with the least hippuric acid release (6.97 ± 0.274 µg/mL). Alcalase, being a serine endopeptidase, had extensively hydrolysed the BBPH thus liberated higher concentration of smaller peptides which were otherwise buried deep the native protein. This rationale can justify the increasingly better ACE inhibitory activity of the time-gradient BBPH hydrolysates. The IC_50_ values of the BBPHs were also mentioned in the figure 1. BBPH_A120_ has shown the least IC_50_ value, which again substantiated the ACE inhibitory activity of BBPH_A120_ as inferred from the inhibition of hippuric acid release.

The results were in agreement with recent studies that mentioned that alcalase mediated hydrolysis of food proteins had produced bioactive peptides with higher ACE inhibitory potentials (Chen, Wang, Zhong, Wu, & Xia, 2012; Forghani et al., 2016), which was inferred to be due to the endopeptidase activity of alcalase.

### 3.2. Identification of the peptide sequence

Based on the degree of hydrolysis and ACE inhibition activity, <3kDa fraction of BBPH_A120_ was analyzed by MALDI-ToF mass spectrometry to identify the resultant peptides due to the alcalase hydrolysis. From the mass-spectrum, five peptides (molecular weights ranging from 914.608 Da to 1653.991 Da) had been identified with significantly high intensity (> 1 × 10^4^a.u.) compared to other peptides (figure 2). *De novo* sequencing of these small peptides revealed high concentration of proline, along with other hydrophobic amino acids (**supplementary table 1**). Current research on ACE inhibitory peptides accommodates quantitative structure-activity relationship (QSAR) approach for initial screening of the peptide sequences (Majumder & Wu, 2010; Nongonierma & FitzGerald, 2016). Previous reports showed that presence of hydrophobic amino acids in the C-terminal was the key feature that influenced the ACE inhibitory action of the peptides and also their binding to the inhibitor binding site in the close proximity of the zinc molecule within the quaternary structure of human ACE (Nongonierma & FitzGerald, 2016; Castellano et al., 2013). Based on these observations, the hydrophobic amino acid rich peptide sequences were checked for overlapping sequences from AHTPDB (http://crdd.osdd.net/raghava/ahtpdb/), a comprehensive database of experimentally validated antihypertensive peptides. The peptide sequences identified from the mass spectrum of ultrafiltrate fraction of BBPH_A120_, had shown significant sequence overlap with many previously reported ACE inhibitory peptide sequences that had shown significantly high *in vitro* ACE inhibitory activity, as demonstrated by the low IC_50_ value (**Supplementary table 1**).

**FIGURE 2:**
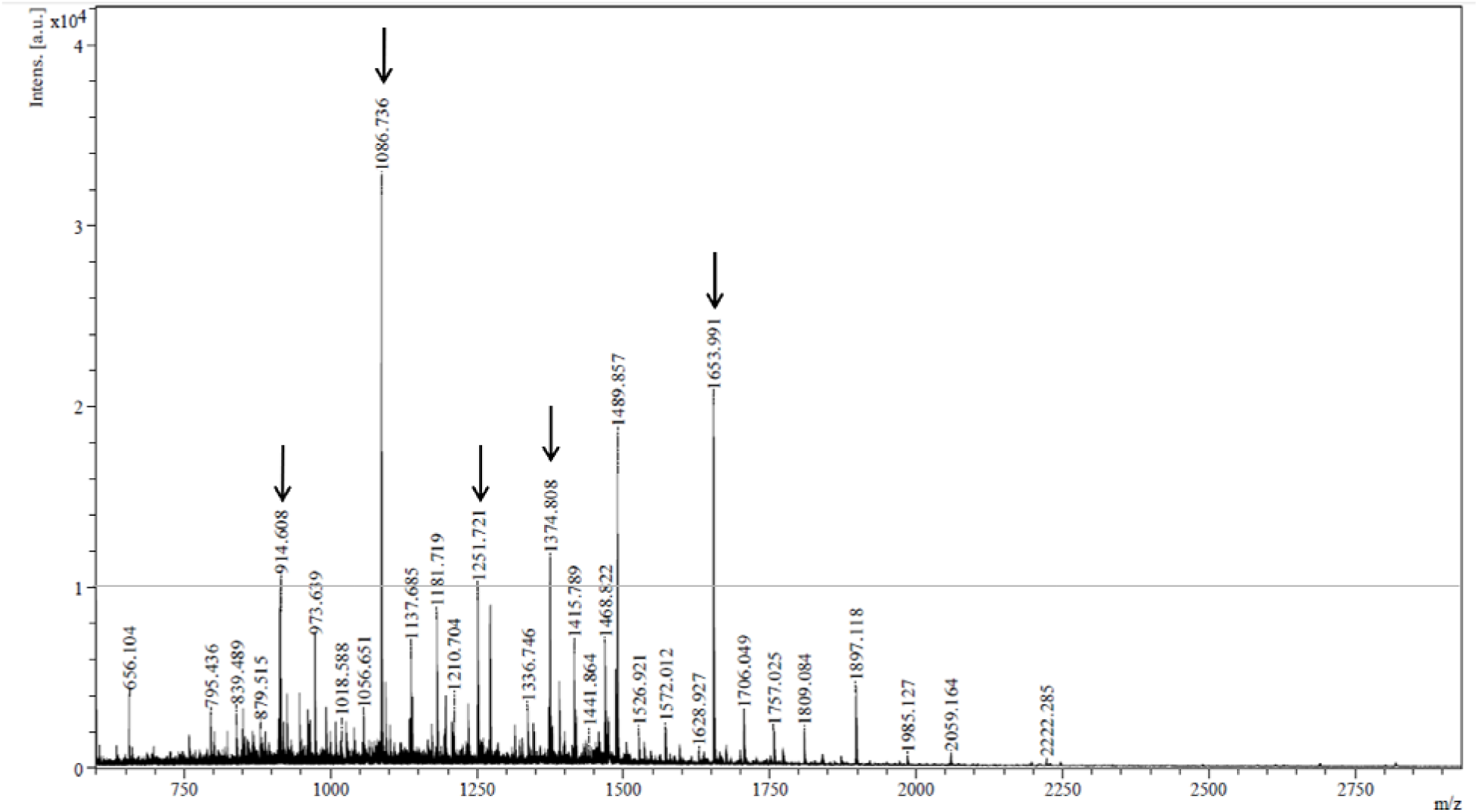
MALDI-TOF SPECTRUM OF THE *B. BENGALENSIS* PROTEIN HYDROLYSATE OF ALCALASE AT 120 MINUTES. THE ARROWS MARK THE PEPTIDES SELECTED FOR THE *DE NOVO* SEQUENCING. THE RED LINE ACROSS SPECTRUM MARKS THE BASELINE INTENTSITY OF 1 × 10^4^ A.U.

Interestingly Belpep which was to be the most concentrated peptide in the BBPH_A120_ mass spectrum (3.25 × 10^4^ a.u.) was also found to have highest sequence overlap with the previously mentioned AHTPDB database. Based on these observations, pure Belpep sequence (IIAPTPVPAAH) was synthetically prepared in order to analyze the molecular mode of inhibition. The purified Belpep had shown a IC_50_ value of 8.52 ± 0.779 µg/mL which reflects strong inhibition on proteolytic activity of ACE that release the hippuric acid from hippuric acid-histidine-lysine (HHL) (figure 1). The C-terminal sequence of Belpep was highly hydrophobic that might have facilitated its entry within the inhibitor binding pocket and resulted into its efficient ACE-inhibitory activity (Castellano et al., 2013).

### 3.3. Inhibitory Kinetics Study

These observations lead to the mechanistic assessment of the mode of belpep mediated inhibition of the ACE, which were elucidated by the Lineweaver-Burk plot as depicted in figure 3, which showed the kinetics of the Enzyme-substrate (ES) reaction in presence or absence of Belpep. From the Lineweaver-Burk plot (figure 3), it is evident that the mode of inhibition by the Belpep was of typical uncompetitive type. In uncompetitive inhibition, the inhibitor doesn’t have any affinity towards the catalytic site of the ACE. Rather the inhibitor preferentially binds to the ES complex, thereby hindering the release of the product and the enzyme from the ES complex. In absence of product, the substrate affinity of the ACE gets increased, as suggested by obvious decrease in K_m_ value in comparison to the same from the uninhibited ES reaction (Table 1). Also, the V_0_ and V_max_ of the belpep inhibited reaction were found to be decreased compared to the control uninhibited ES reaction. In fact the Lineweaver-Burk plots of the control and the belpep inhibited ES reactions showed no sign of convergence, neither at the y-intercept nor at the x-intercept (figure 3), which was characteristic of the uncompetitive inhibition of ES reaction (Jao, Huang, & Hsu, 2016; Forghani et al., 2016).

**Figure 3:**
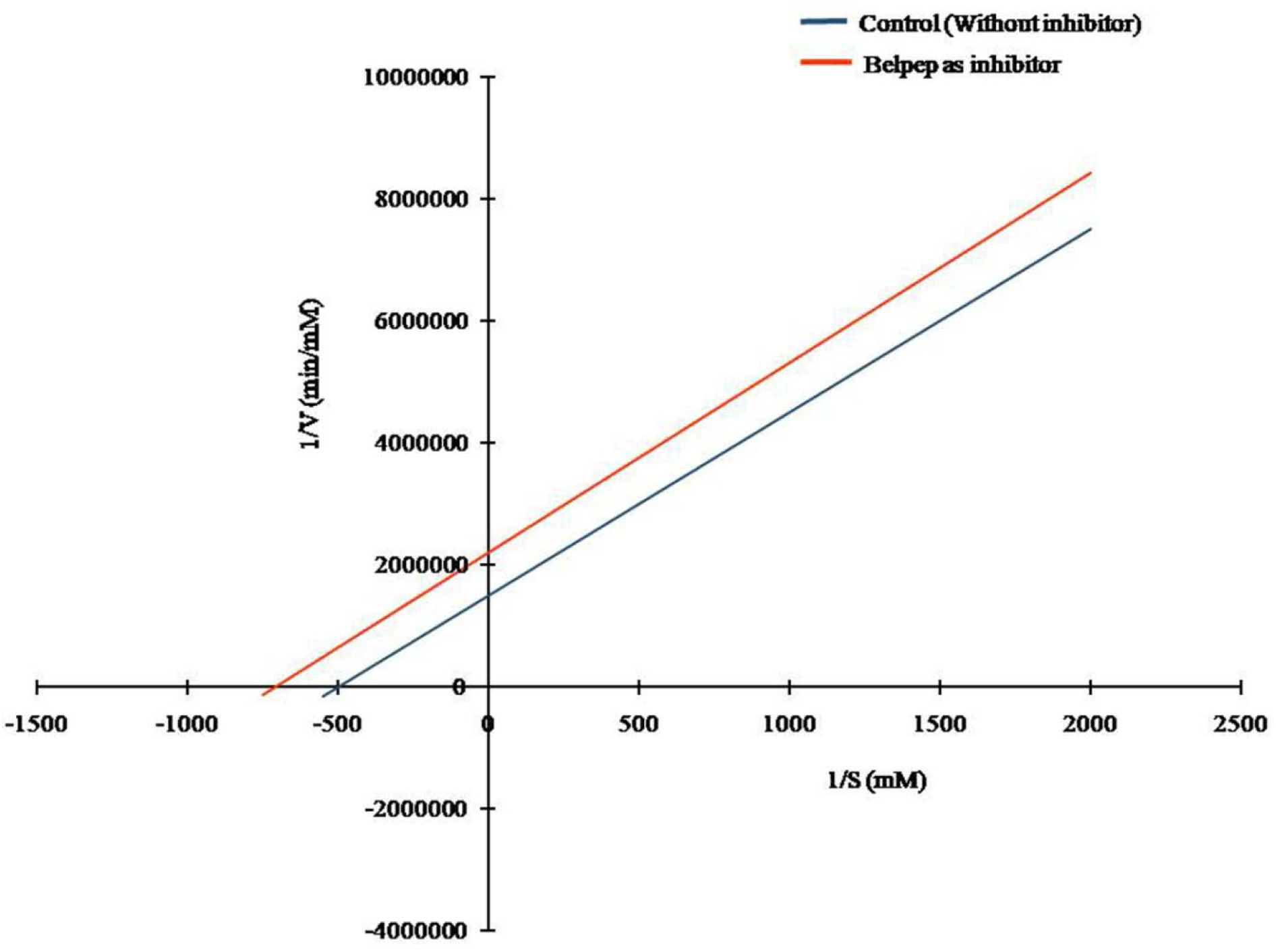
ACE inhibitory kinetics of the peptide isolated from ultrafiltered 3kDa permeates of *Bellamya bengalensis* protein alcalase hydrolysate. Lineweaver –Burk plot of ACE inhibition by the peptide isolated from ultrafiltered BBPH_A120_. The ACE inhibitory properties were evaluated both in presence and absence of inhibitor. 1/[S] and 1/V represent the reciprocal substrate concentration and velocity of the reaction, respectively.

**Table 1:**
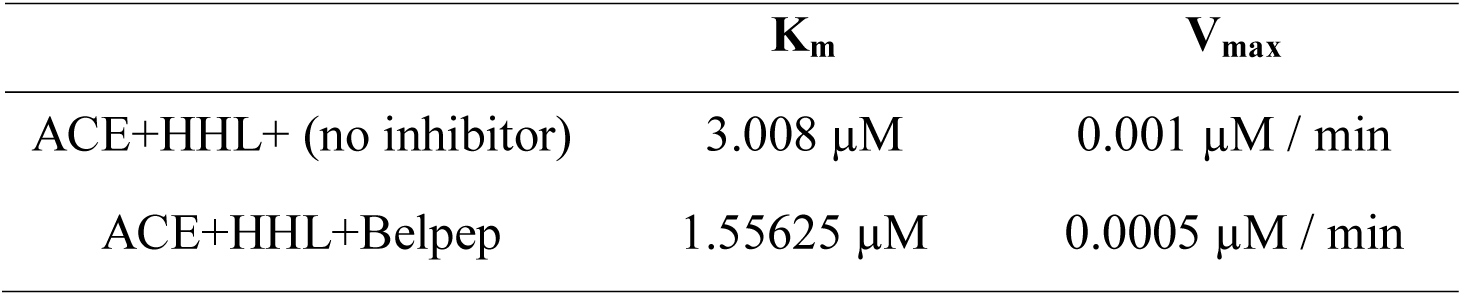
ACE inhibitory kinetics parameters of the peptide isolated from ultrafiltered BBPH_A120_ (*Bellamya bengalensis* alcalase hydrolysate at 120min).

### 3.4. Isothermal Titration Calorimetry

ITC experiments were conducted to confirm the mode of inhibition of ACE by Belpep based on the thermodynamic parameters of its binding to the ACE and HHL. The thermograms and binding isotherms of the Enzyme-Substrate titration at optimal conditions in absence and presence of the belpep were presented in figure 4.

**Figure 4:**
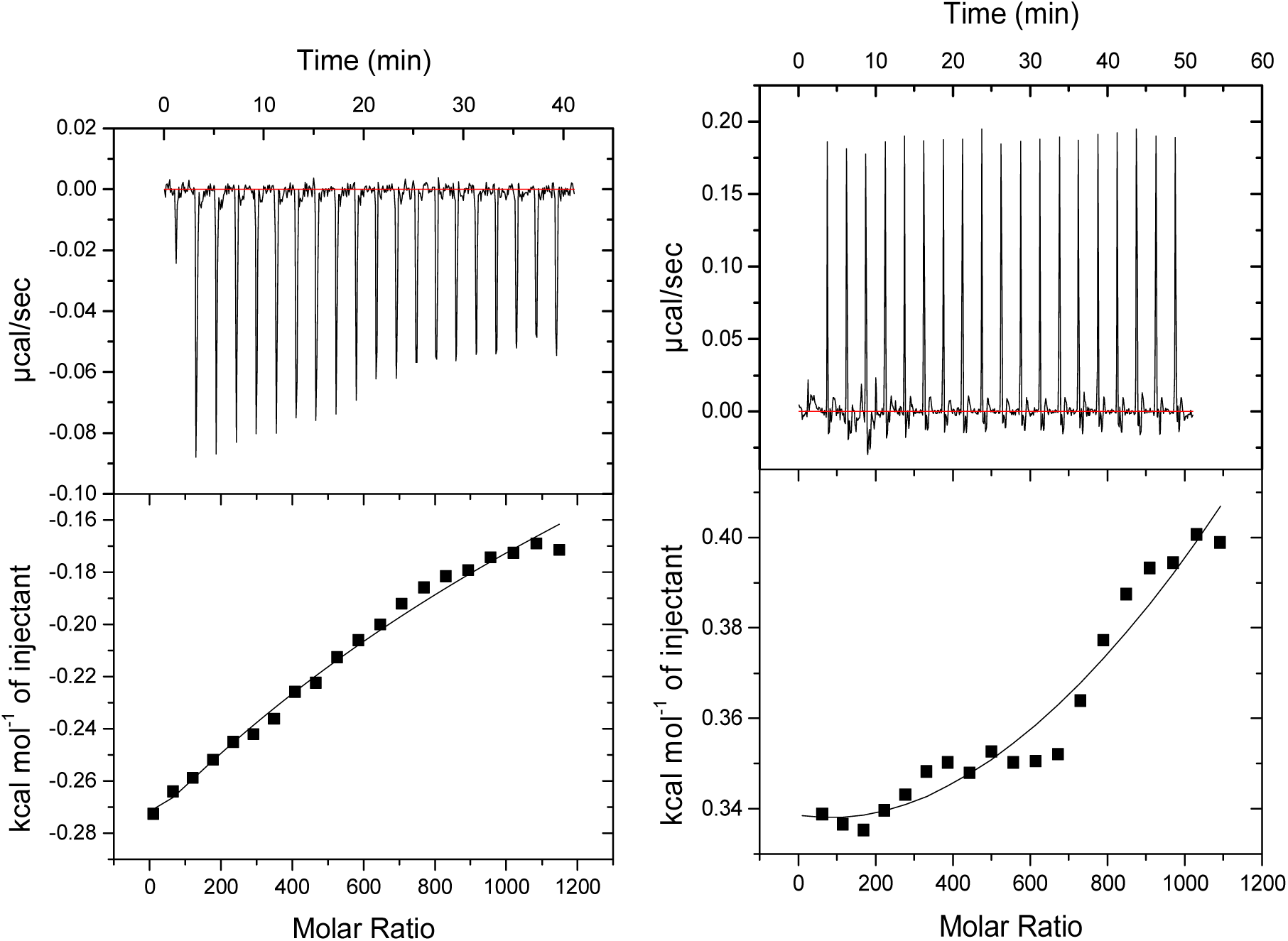
The isothermal titration calorimetric analysis of the binding of angiotensin converting enzyme (ACE) at a concentration of 357nm to the substrate HHL at a concentration of 1mM and without any inhibitor (Figure 4A); or with belpep as inhibitor at a concentration of 0.5mM (Figure 4B). The upper panel shows a typical ITC curve, showing the heat release as a function of time depicted in the thermogram for binding of the ligand and the enzyme at pH 8.3 and temperature 37°C whereas the lower panel of the figure shows the isotherm depicting the integrated heat evolved per mole of the inhibitor for addition of each injection with respect to the total molar ratio of the ligand over the enzyme concentration.

Figure 4A showed a typical titration curve for the binding of HHL to ACE under optimal condition as ACE hydrolyze hippuric acid from HHL via an exothermic reaction (enthalpy change −5.66×10^6^ kJ/mol) (Ni et al., 2012). As the ACE get titrated with substrate (HHL), the catalytic sites were progressively get saturated and as a result the reaction tended towards equilibrium, net heat release diminished and only background heat of dilution peaks were remained as shown in the thermogram. At the end of reaction, it tended towards achieving saturation and it caused the enthalpy change due to the addition of moles of ligand to the enzyme solution at isothermal condition.

But in presence of an inhibitor belpep, the exothermic nature of the ACE-HHL reaction was changed to an endothermic pattern (figure 4B). Previous reports also had shown similar endothermic pattern of thermogram when standard inhibitor like lisinopril was used to inhibit the ACE-HHL reaction (Andújar-Sánchez, Cámara-Artigas, & Jara-Pérez, 2004). The area under each small injection peaks (heat absorbed per injection) in belpep inhibited ES reaction signified very small rate of heat change (0 to 0.20 µcal/sec) in comparison to the uninhibited ES reaction, which was evident from the corresponding binding isotherms as well.

Interestingly, the binding isotherm curvature of the belpep inhibited ES reaction indicated towards a 2 site co-operative ligand binding to the ACE molecules (Freiburger, Auclair, & Mittermaier, 2009; Brautigam, 2015). This observation was coherent to the uncompetitive inhibitory nature of Belpep. Initial slow but gradual increase in the rate of heat change with increase in the molar concentration of the ligand indicated towards initial substrate binding to the ACE molecule, which in turn promoted the binding of belpep to a secondary binding site on the ACE as confirmed by a comparatively sharp increase in the rate of secondary heat change. Thus, ITC experiments validated the binding of the belpep to the ACE imparting inhibitory effect, coherent with the kinetics of enzyme inhibition. These observations were further validated by the molecular docking of the inhibitor belpep on to the ACE molecule.

### 3.5. Molecular Docking

The automated molecular docking pose involving the binding of the belpep to the ACE, with minimal binding energy was depicted in figure 5A. The hydrophobic nature of the belpep was evident from its sequence. Therefore the binding site of Belpep should be logically located within the core of the enzyme molecule, which was indeed what was found in the docked structure. Yet there were considerable differences between the amino acid residues that were found to be actively interacting with the Belpep, and the same that were involved in hydrogen bonding with lisinopril (Natesh et al., 2003). Belpep was found to be involved in hydrogen bonding with amino acid resides Asp376, His344, Tyr480, Glu123, Gln242, Lys 468 and Phe469, which were not part of the S_1,_ S_1_ and S_2_ substrate binding pockets of ACE that were occupied by lisinopril within the co-crystal (PDB no. 1O86) (Natesh et al., 2003) (figure 5B). Binding of substrate to this core groove of ACE within the close proximity of Zn^2+^ ion, probably rearranged the groove a little bit, so that a second ligand binding locus was formed about 4-5 Å away from the active site Zn^2+^ ion. This secondary binding locus was acquired by the Belpep to affect its uncompetitive inhibitory activity, which was coherent to the enzyme kinetic data. No direct interaction between the Belpep and the Zn^2+^ ion of ACE (Figure 5A) was observed in this simulation, although such direct interaction of the ligand with the Zn(II) atom of ACE for exerting strong inhibitory effect was a subject of controversy among the pertinent literature (Wu et al., 2015; Rawendra, Chang, Chen, Huang, & Hsu, 2013; Jimsheena & Gowda, 2010; Guo, Pan, & Tanokura, 2009).

**Figure 5:**
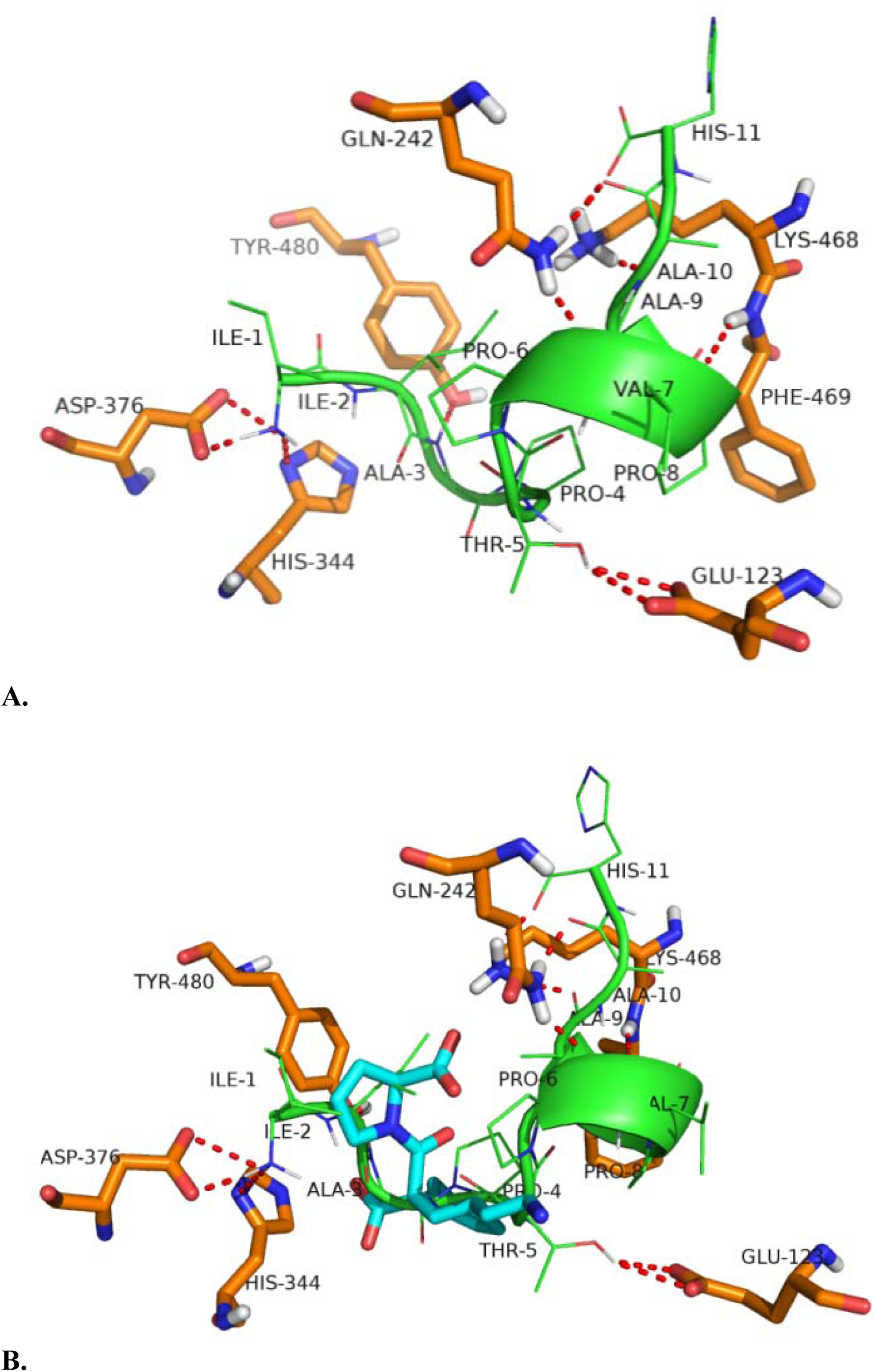
Docking of Belpep (IIAPTPVPAAH) with ACE shows that it binds at the core of the protein (Figure 5A). Superposition of docked structures with PDB 1O86 showed that it binds to similar locus compared to lisinopril (Figure 5B).

## 4. Conclusion

The present study confirmed the presence of ACE inhibitory peptides as extracted from the hydrolysates of protein concentrate of *Bellamya bengalensis*. The enzymatic hydrolysis yields low molecular weight peptides with extraordinary properties like lowering hypertension by means of ACE inhibition. This convenient mode of preparation of hydrolysate fraction produced sufficiently potent ACE inhibitors of commercial relevance which can act as a value-added substance in food formulations. The inhibitory mechanism perceptions with the combination of ITC and molecular docking designate a novel approach towards the *in vitro* inhibitory mechanism study. Bioactive peptide extraction from the meat of *Bellamya bengalensis* facilitates the opening up of innovative economic opportunities which can amplify the utilization of *Bellamya bengalensis* meat.

## 6. Acknowledgement

The authors acknowledge Dept. of Biochemistry, IPLS, Centre for nanoscience and nanotechnology of University of Calcutta for the infrastructural support. Financial support provided by Ministry of Food Processing, Government of India is also duely acknowledged.

