## supplementary figure 1 and table 1 for "*In vitro* and *in silico* analyses of the angiotensin-I converting enzyme inhibitory activity of peptides identified from *Bellamya bengalensis* protein hydrolysates"

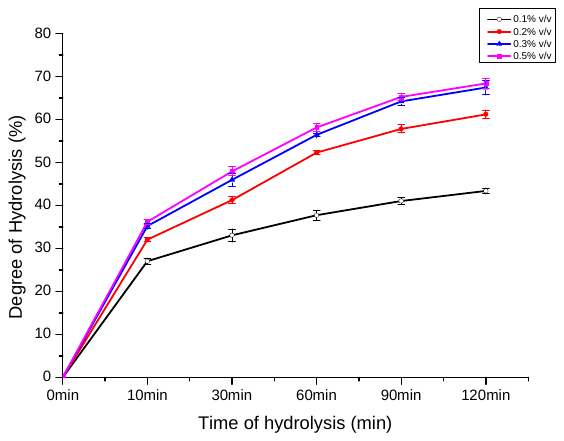

Supplementary Figure 1: Degree of hydrolysis of *B. Bengalensis* protein isolate by alcalase with respect to time.

**Supplementary Table 1:** Sequence-matching in relation to documented ACE-inhibitory peptides.

| **Parent Mass (Da)** | **Sequence** | **ACE inhibitory peptide from AHTPDB database** | **IC_50_ value** | **Source** | **Reference** |
| --- | --- | --- | --- | --- | --- |
| 914.608 | L**TPV**PGSPF | LVNDLV**TPV**FDNL | 4.00 mg/ml | Fungi (Mushroom) | Kang et al., 2013. |
|  |  | **TPV**VVPPFLQP | 749 μM | Cheese whey protein | Puchalska, Marina Alegre, & García López, 2015. |
| 1086.736 | IIA**PTPVP**AAH | **PTPVP** | 256.41 μM |  | Sagardia, Roa-Ureta. & Bald, 2013 |
|  |  | **PTPVP** | >1000 μM |  | Puchalska, Marina Alegre, & García López, 2015. |
|  |  | **PTPVP** | 256.41 μM | *Pork meat* | Escudero, Sentandreu, Arihara, & Toldrá, 2010 |
| 1251.721 | TI**GAP**DGIPSAPR | GPA**GAP**GAA | 37.15 μM |  | Sagardia, Roa-Ureta. & Bald, 2013 |
| 1374.808 | HEFPG**VVV**GANDD | P**VVV**PPFLQP |  | Milk | Meisel & Schlimme, 1990. |
|  |  | TP**VVV**PPFLQP | 749 μM | Cheese whey protein | Puchalska, Marina Alegre, & García López, 2015. |
|  |  | **VVV**PP | 284.4 μM |  | Sagardia, Roa-Ureta. & Bald, 2013 |
| 1653.991 | **LNP**GAGLPRGPNGADTF | **LNP** | 43 μM |  | Tan et al. 2013. |
|  |  | **LNP**A | 312 μM |  | Tan et al. 2013. |
|  |  | L**LNP** | >1000 μM | *Milk*  *(human β casein)* | Kohmura, Nio, & Ariyoshi, 1990. |
|  |  | **LNP**A | 312 μM | *Alpha-Zein hydrolysate*  *(Zea mays)* | Yano, Suzuki, & Funatsu, 1996. |
